# Jack-of-all-trades paradigm meets long-term data: generalist herbivores are more widespread and locally less abundant

**DOI:** 10.1101/2020.11.27.401778

**Authors:** Chanchanok Sudta, Danielle M. Salcido, Matthew L. Forister, Thomas R. Walla, Santiago Villamarín-Cortez, Lee A. Dyer

**Affiliations:** University of Nevada, Reno; University of Nevada; Colorado Mesa University

**Keywords:** Diet breadth, diet specialization, abundance, plant-herbivore, scaling, tropics, long-term data, generalists, specialists

## Abstract

Ecological specialization is one of the most interesting and perplexing attributes of biological systems. While certain macroecological patterns, such as an increase in specialization at lower latitudes, have long been subjects of investigation, there is much yet to be learned about inter-specific variation in specialization within diverse communities. High levels of specialization have been documented for some dominant ecological interactions, such as parasitism and herbivory, but much less is known about the relative abundance of specialists and generalists within those broad functional groups. We examine untested assumptions about the positive association between local abundance and dietary specialization using a 17-year dataset of caterpillar-plant interactions in Ecuador. Our long-term data consist of experimental verification of caterpillar-plant associations and include standardized plot-based samples as well as general, regional collections of caterpillars, allowing for investigations across spatial scales and using different indices of abundance for 1,917 morphospecies of Lepidoptera (“caterpillars”) from 33 families. We find that more specialized caterpillars are locally more abundant than generalists, consistent with a key component of the “jack of all trades, master of none” hypothesis, which has otherwise received poor to mixed support from previous studies that have mostly involved fewer species and shorter time series. At larger scales, generalists achieve greater prevalence across the landscape, and we find some evidence for geographic variation in the abundance-diet breadth relationship, in particular among elevational bands. Interspecific variation in abundance had a negative relationship with diet breadth, with specialists having more variable abundances across species. The interesting result that more specialized species can be both rare and common highlights the ecological complexity of specialization.

**Statement of authorship:** CS wrote the first draft of the manuscript. DMS, MLF and LAD contributed substantially to consequent drafts and revisions. LAD, TRW, SV and DMS collected field data. CS, MLF and LAD generated research questions and designed statistical analyses. CS, DMS, LAD and MLF conducted statistical analyses.

**Data statement:** Data supporting the results and conclusions can be found on a website caterpillars.org. Statistical analyses supporting the results are available upon request.

## Introduction

Insects are a remarkably diverse taxon, and all ecological guilds of insects engage in some of the most consequential and common ecological interactions. The fields of ecology and evolutionary biology have benefited from studies of insect interactions, especially studies on niche specialization, which has attracted decades of attention (Ehrlich & Raven 1964; Mani 1968; Forister *et al.* 2012). A common expectation for specialization has been that “a jack of all trades is a master of none,” which is, in the ecological context of dietary specialization, the idea that a generalist might be able to consume a wide range of resources but will not be particularly well adapted to any one of them (MacArthur 1972; Futuyma & Moreno 1988). Two predictions are commonly associated with this idea. First, functional trade-offs have long been hypothesized as the most parsimonious explanation for the predominance of specialists. If adaptation to one resource comes at the expense of lower performance on another resource, then specialists will be common, and generalists will be favored only under certain circumstances. For example, a single-locus antagonistic pleiotropy for the ability to digest different food types favors specialization on one food type, whereas generalists might only be favored under conditions of fluctuating host plant availability. Functional trade-offs in dietary specialization have received only minimal support (Hardy *et al.* 2020), and are not our focus here. We are interested in a second prediction involving ecological consequences of specialization, specifically the idea that specialists should be locally more abundant, while generalists might never be as abundant in any one location but might occupy more places across the landscape. This prediction has often been stated in experimental terms for a small number of taxa (e.g., specialists will outperform generalists when tested on a common resource), which highlights the lack of relevant ecological, field-based datasets in the literature (Dyer *et al.* 2015).

Studies that close the knowledge gap with respect to diet breadth and abundance at different scales are likely to contribute substantially to our understanding of specialization and the ecology and evolution of plant-insect interactions. There is no single measure of specialization that captures the subtleties of insect diet breadth (Table S1), thus we have taken a multifaceted approach that includes taxonomic and phylogenetic diet breadth as well as a complementary, ordinated index based on the community of observed plant-herbivore associations (Table S1). We use a 17-year empirical dataset from Ecuador, which included replicated elevational gradients, to examine associations between diet breadth and caterpillar (larvae of Lepidoptera) abundance, and to include potential effects of spatial scale on these relationships. With these empirical data, we ask the simple question of whether there is an association between abundance and diet breadth and how that relationship might vary across different scales of observation. We hypothesized that caterpillar abundance would decrease with increasing diet breadth, consistent with the “jack of all trades is a master of none” hypothesis, especially at smaller spatial scales where the local demographic advantage of being well-adapted to a host plant should be evident. In contrast, neutral models predict positive associations between diet breadth and abundance, especially at the landscape level (Forister & Jenkins 2017). Consistent with this, at larger spatial scales, we expected to discover an advantage to dietary generalization: because generalists can be found on multiple host species, they should turn up in more surveys and plots, reflecting broader landscape-level occupancy.

## Methods

### Study location & collection protocol

We used 17-years of host plant-caterpillar interaction data collected within the Eastern Andes of Ecuador between 2001–2018 (Fig. 1a; 420 km^2^ from 00.36° S and 77.53° W) along an elevational gradient (100-3800m) that included low montane forests and paramo habitat to explore relationships between caterpillar diet breadth and density. Caterpillars were found by visual inspection of leaves using two methods: general collecting and sampling within standardized circular plots (10 m diameter). Both collection protocols and rearing methods are fully described in previous publications (Dyer *et al.* 2007; Salcido *et al.* 2020). Briefly, general collection consisted of opportunistic sampling of caterpillars along roads, trails, transects, or off-trail searches while plots were located at least 10 m from roads or trails and included estimates of plant diversity and genera-level leaf area. For both methods, caterpillars were collected and assigned a specific record number and reared to adulthood. Caterpillars and their host plants were identified to the lowest possible taxonomic level. In cases where the resolution of taxonomic identifications was limited to family or genus, caterpillar or host plant morphotypes were assigned based on family or genus designations in combination with morphological descriptors. We removed data for which either host plant or caterpillar family and descriptors were unavailable.

**Figure 1.**
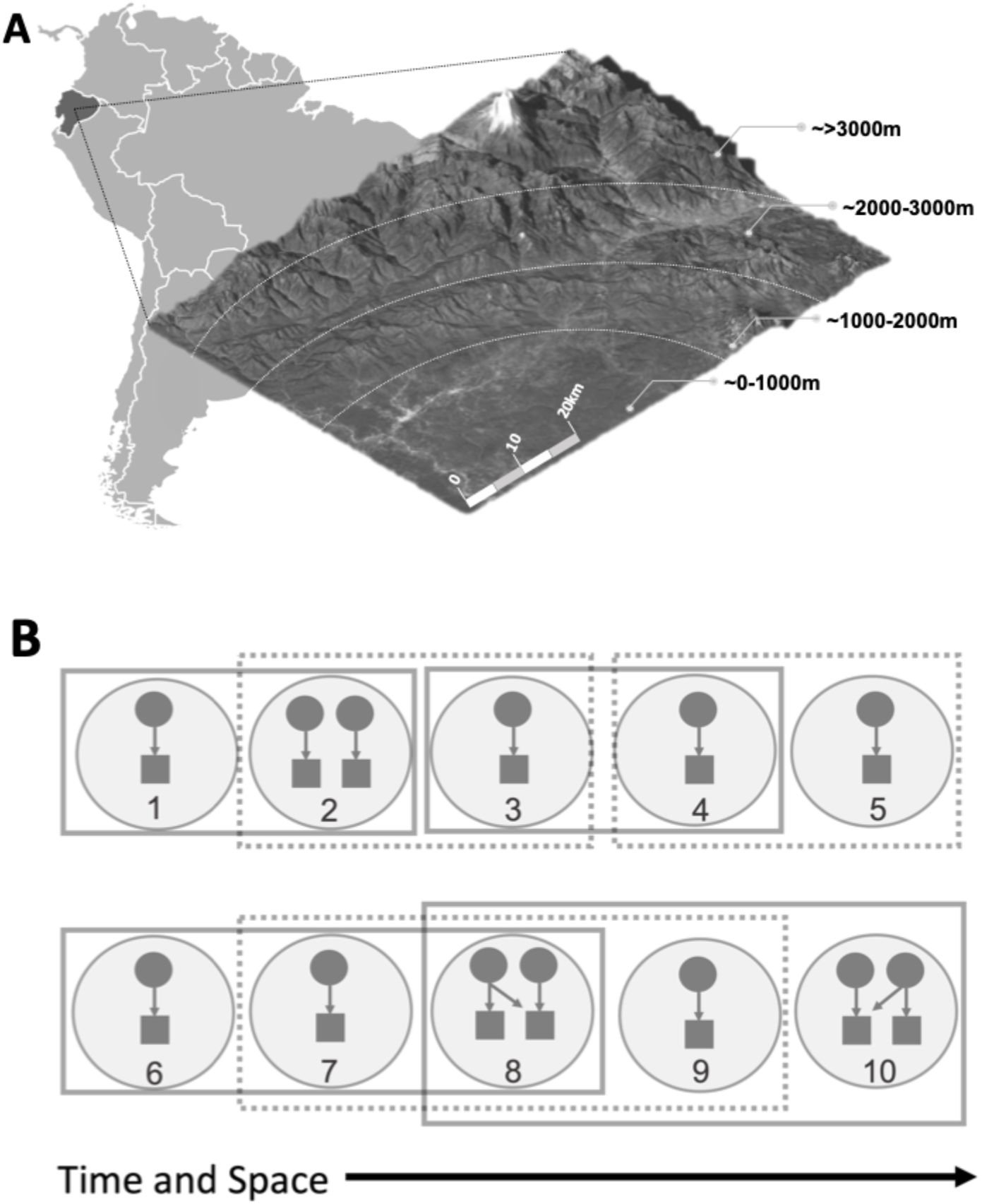
This study took place in the Northeastern Andes in Ecuador, and plot-level data were aggregated to examine the effect of increased scale on the relationship between density and diet breadth. A) Samples were collected within the Napo Province across different elevations. Plots were categorized into one of four distinct elevational bands (0-1000m, 1001-2000m, 2001-3000m, >3000m from sea level); although substantial species turnover occurs at finer scales, we only had sufficient data for these groups. Lines delineating the different elevations are not exact and are intended to be used as a heuristic tool. B) Effects of scale were evaluated at the plot level by sequentially aggregating plots (circles) using a moving window method (solid and dashed boxes). The moving window method was applied to the plot data iteratively: each time the size of the window (the number of plots aggregated) increased by one more plot. For a window size of five (e.g., plots 1-5), every two plots were sequentially aggregated. In the next iteration, every six plots were sequentially aggregated (e.g., plots 1-6). For each aggregate and caterpillar species (circular nodes), abundance was summarized. Density was calculated based on the combined area or corrected for the leaf area of associated host plant genera (square nodes). Density across all plot aggregates for a given species were averaged.

### Diet breadth and density estimates

We used host plant and caterpillar taxonomic identifications from both general collections and plots to calculate diet breadth of each caterpillar species. For our analyses, taxonomic diet breadth is the total number of unique host plant species consumed by a given caterpillar species or morphotype using all data from general collections and plots. We repeated all analyses using complementary methods for ordinated diet breadth and phylogenetic diet breadth (results are reported in detail in the supplementary materials). Ordinated diet breadth is based on all observed associations between plants and caterpillars and involves calculating multivariate distances among hosts in ordination space that is structured by the plant-caterpillar associations (Fordyce *et al.* 2016). For example, a caterpillar associated with three hosts that are rarely used by other herbivores will have a broader ordinated diet breadth than a species that uses three hosts that are frequently used in combination by other caterpillar species. Thus, this method potentially encompasses functional similarities (e.g., shared plant chemistry) among suites of hosts (Fordyce *et al.* 2016). Phylogenetic diet breadth is measured as a phylogenetic distance among all host plants of a given herbivore species (Symons & Beccaloni 1999; Novotny *et al.* 2002; Ødegaard *et al.* 2005; Forister *et al.* 2015). Both ordinated diet breadth and phylogenetic diet breadth were calculated at the host plant family level. We are aware that other and often more complex phylogenetic indices of diet breadth exist (Jorge *et al.* 2017), but we have used the index most easily interpreted (involving branch lengths spanning hosts for a particular herbivore) for comparison with non-phylogenetic indices. We generated phylogenetic distances among host plant families using the R package picante (as in Forister *et al.* 2015) and an angiosperm phylogeny from Davies *et al.* (2004).

In contrast to diet breadth which used all available information, we only used data collected within plots to estimate caterpillar densities and landscape occupancy at plot and elevation scales. We calculated three indices of abundance or occupancy. First, the abundance of each caterpillar species was summed across 10-m diameter plots (78.5 m^2^) and standardized by the density of relevant host plant within the same plots. Second, we calculated an index of landscape occupancy as the frequency of observation for a particular caterpillar species across plots. For these first two measures, calculations were made for plot and elevation scales for which adequate data were available. Third, we explored an alternative measure of density by calculating density per unit plot area and reported associated results in the supplement.

### Spatial scales

To evaluate the effect of spatial scale on associations between diet breadth and abundance we examined patterns at various scales (in order of increasing area): 1) plot scales, being individual plots as well as those that were aggregated to form larger areas, 2) discrete elevational bands, and 3) a regional scale encompassing our entire study area. For the plot scale, data were collected from a total of 455 plots across the study area. We examined the effect of scale on the relationship between diet breadth and density at the plot-level by aggregating plots into temporally and spatially-adjacent groups then calculating the average density and occupancy for each caterpillar species across all plot groups. This process was iterated for plot groupings of increasing size. To group plots, a moving window method was applied (Fig. 1b). The window size was equivalent to the number of plots grouped together (*n* = 2 - 455 plots) and moved sequentially (in order of time that the plots were completed) along the plot data in order to maximize spatial and temporal similarities among plots. For each iteration, the window size increased by one plot. For example, for a window size of two, caterpillar abundance and leaf area from plot 1 and plot 2 were, summarized followed by all iterations of plot *n*-1 and plot *n*. In the subsequent iteration, the window size increased to three plots and so on until the window size equaled the number of all plots. Results presented in the main text are limited to groupings of 5 (∼393 m^2^ in total across aggregated plots), 25 (∼1963 m^2^), 50 (∼3927 m^2^), and 250 (∼19535 m^2)^) plots; additionally, results for all possible aggregates were calculated and are presented in the supplement (Fig. S1-S2).

Plots within four distinct elevational bands: 1) 0 m - 1000 m, 2) 1001 m - 2000 m, 3) 2001 m - 3000 m, and 4) above 3000 m were aggregated (Fig. 1a); these elevational bands are ecologically distinct (Pyrcz *et al.* 2009). Density was calculated based on the total caterpillar abundance within each elevational band per available host plant area. While turnover is substantive at a finer elevational scale (e.g., Sklenár & Ramsay 2001), elevational bands were defined based on data availability and to allow for sufficient data at distinct elevations.

Finally, to examine caterpillar abundance at the regional level, total abundance for each caterpillar species was summarized across the entire dataset. This included caterpillars collected from both plot-based and opportunistic collection methods. For regional data, species with more generalized host ranges can be observed more often simply because they occur on a wider range of host plants (and are thus encountered more frequently), so we calculated the number of times that a caterpillar has been collected, divided by the number of hosts on which it occurs (in other words, average abundance per host). For example, if a particular caterpillar species has a total abundance of 10 and a diet breadth of 2 (it uses 2 host plant species), the density per host species is 5 at this region scale. Prior to the analyses described here, we identified and removed extreme outliers and singletons (species for which there was only one record in the dataset). Due to its unusually high abundance from sampling bias associated with related research projects, we also removed *Eois olivacea* (Geometridae: Larentiinae).

### Statistical inference

We used Bayesian linear models to estimate beta coefficients for the association between abundance or occupancy and diet breadth for plot, elevational and regional scales. Models were fit in JAGS (version 3.2.0) utilizing the “rjags” package in R (R version 3.6.3) using (for each analysis) two Markov chains and 1,000,000 steps each; performance was assessed through examination of chain histories (burn-in was not required because of rapid convergence), effective sample sizes and the Gelman and Rubin convergence diagnostic. Response variables (abundance and occupancy) were modeled with assumptions of normally distributed residuals with means dependent on an intercept plus diet breadth as a predictor variable. We used weakly informative priors: priors on beta coefficients were normal distributions with mean of zero and precision of 0.001 (variance = 1000); priors on precisions were modeled as gamma distributions with rate = 0.1 and shape = 0.1. We report medians from posterior distributions for beta coefficients and corresponding 95% credible intervals (CI). Values for density, occupancy, and taxonomic diet breadth were log transformed (log_10_) prior to analyses.

## Results

Caterpillar density per available host area declined with increasing diet breadth at the level of individual plots (Fig. 2a, Table 1, Fig. S1). The overall relationship between diet breadth and caterpillar density per available host area was also negative: the beta coefficient estimated from linear models was −0.36 (95% credible intervals: −0.48 to −0.23) for all plot clusters (Table 1, Fig. S1), and this negative relationship is not dependent on the number of plots being aggregated. The raw effect size associated with this relationship (predicted from a simple linear regression of untransformed density on diet breadth) can be interpreted as an increase in diet breadth from 1 to 10 host plants yielding a 36% (± 1.2%) decrease in density per leaf area, and this effect size is similar for the other negative beta coefficients reported in Table 1. Similarly, all elevational bands showed a negative relationship between caterpillar density per available host plant area and diet breadth except at the lowest elevation band (0 m −1000 m, Fig. 2b, Table 1) with the same magnitude of effect sizes. Overall, this negative relationship between abundance and diet breadth reflects a tendency for more specialized caterpillar species to be more locally abundant relative to species that consume a greater number of hosts.

**Figure 2.**
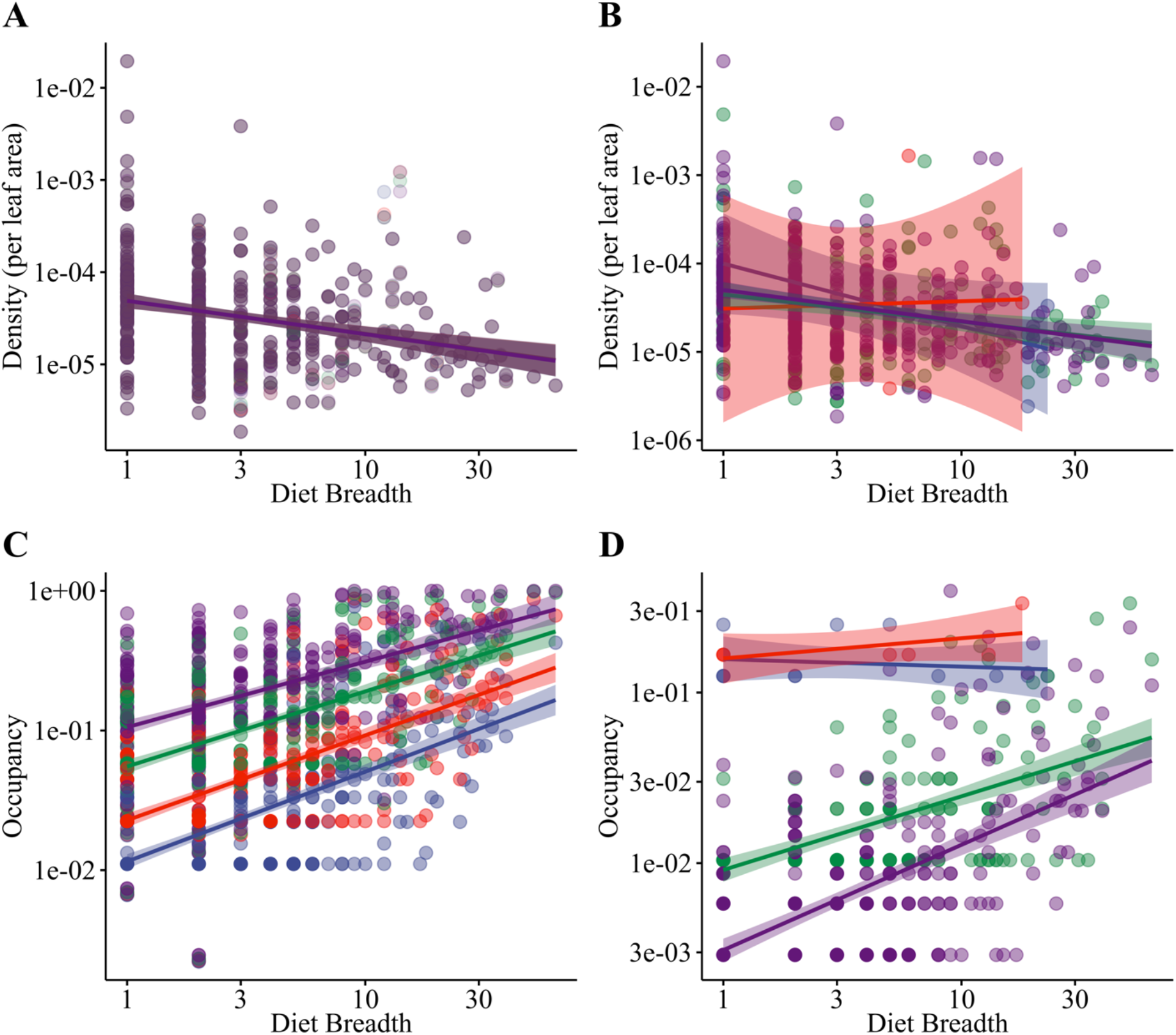
Relationship among diet breadth and caterpillar density corrected for available host area (cm^2^) and occupancy (landscape presence) at different scales of observation: a) and c) depict plot scales, and b) and d) depict elevation scales. Plot-level data were combined for each caterpillar species (points) for various plot aggregates (blue: 5 plots, red: 25, green:50 and purple:250) to evaluate the effect of scale. For the elevation analysis, discrete elevation bands included: 0 - 1000 m (red), 1001-2000 m (green), 2001-3000 m (purple), and above 3000 m (blue). Points were set to a transparency value given the quantity of points, such that darker hues indicate a high density of points.

**Table 1.** Point estimates for beta coefficients (bold) and associated 95% credible intervals for relationship between taxonomic diet breadth and density per available host plant area and occupancy at plot and elevation levels. Plot levels include different cluster sizes which represent the number of aggregated 10-m plots.

| Level of Observation | *Measure of Density β-coefficient [95% CI]* |  |
| --- | --- | --- |
|  | Density per available host plant area | Occupancy |
| **Plot level (cluster size)** |  |  |
| 5 | **-0.35** [-0.47, -0.22] | **0.64** [0.56, 0.72] |
| 25 | **-0.36** [-0.49, -0.23] | **0.54** [0.46, 0.61] |
| 50 | **-0.36** [-0.48, -0.23] | **0.47** [0.39, 0.54] |
| 250 | **-0.35** [-0.47, -0.24] | **0.24** [0.15, 0.32] |
| **Elevation level** |  |  |
| 0 m - 1000 m | **0.10** [-1.72, 1.63] | **0.12** [-0.33, 0.50] |
| 1001 m - 2000 m | **-0.31** [-0.49, -0.12] | **0.43** [0.34, 0.53] |
| 2001 m - 3000 m | **-0.35** [-0.48, -0.21] | **0.62** [0.52, 0.71] |
| > 3000 m | **-0.73** [-1.48, 0.068] | **-0.05** [-0.30, 0.20] |

In contrast, occupancy (frequency of occurrence across the landscape) increased with increases in diet breadth for all plot cluster sizes, but the estimate for the slope was dependent on plot cluster size, as it increased with increasing plot cluster size (Fig. 2c, Table 1, Fig. S2). The raw effect size associated with the overall positive relationship (for all plot cluster sizes) can be interpreted as an increase in diet breadth from 1 to 10 host plants yielding a 18% (± 0.5%) increase in density per plot cluster. However, the relationship between occupancy and diet breadth varied across different elevation bands (Fig. 2d, Table 1). A positive relationship was observed at the middle elevation bands (1001-2000 m and 2001-3000 m) while weak positive and negative relationships between occupancy and diet breadth were observed at the low (0 m - 1000 m) and high (>3000 m) elevation bands respectively (Fig. 2d, Table 1). The positive relationship between occupancy and diet breadth provided evidence that more generalized species occupy more places across landscape.

At our largest spatial scale, using all data, including plots and general collections, we detected a much noisier negative relationship between density per available host plant species and diet breadth (β = −0.076, 95% CI [−0.15, −0.0061], Fig. 3). Nevertheless, the raw effect sizes indicate an appreciable decrease in density with diet breadth, with a change from feeding on 1 to 10 host plant species yielding a 29% (± 19%) decrease in density per leaf area. When density was calculated per plot area, the patterns at both plot and elevation scales were similar to trends observed for occupancy (Fig. S3-4, Table S2).

**Figure 3.**
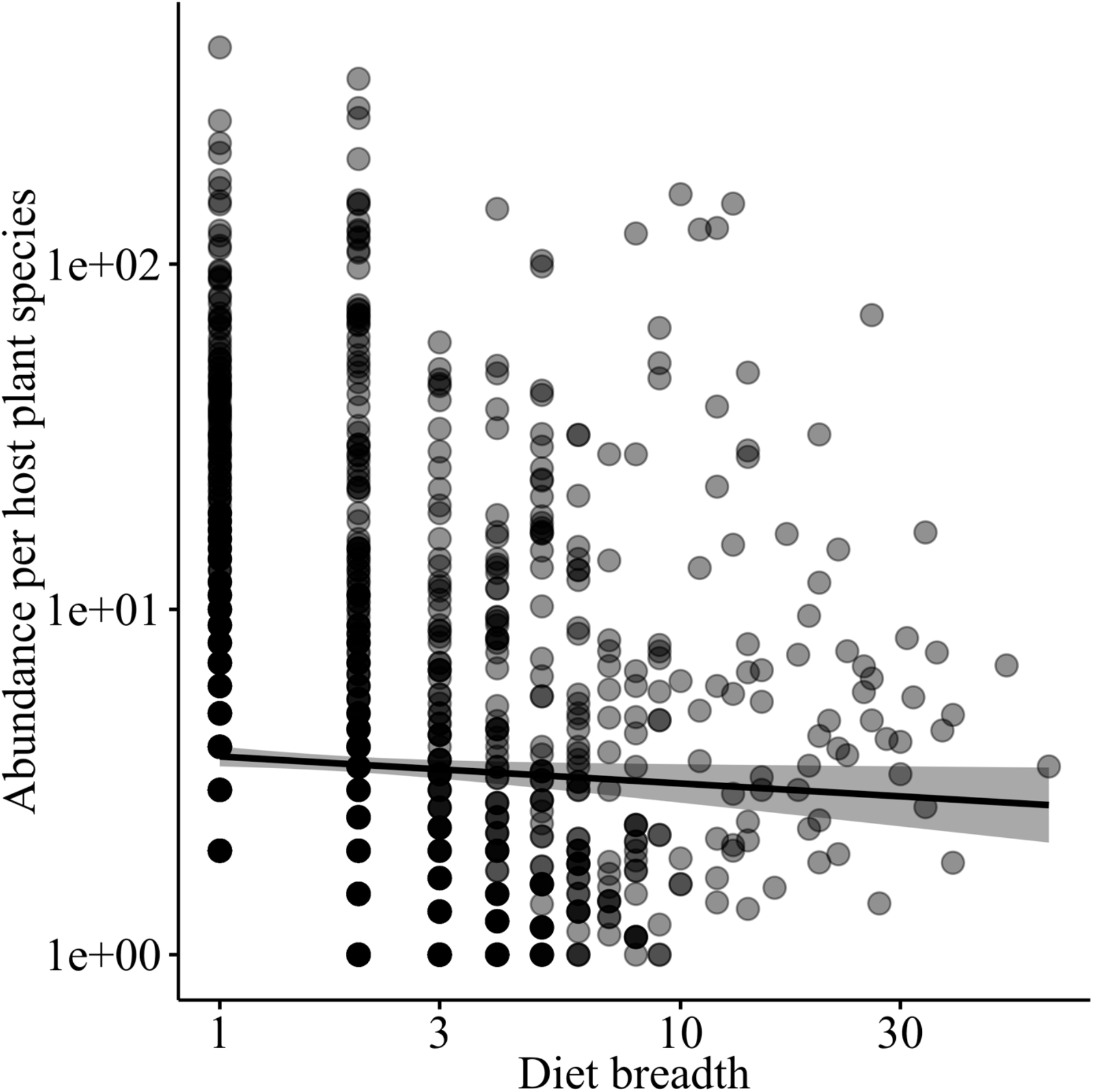
Relationship between caterpillar density per host plant species and diet breadth at a regional scale. Points were set to a transparency value given the quantity of points, such that darker hues indicate a high density of points.

Inferences about associations between diet breadth and abundance were robust to the use of different specialization indices, specifically ordinated diet breadth and phylogenetic diet breadth. Our results using ordinated diet breadth (Fig. S5-8; Table S3) and phylogenetic diet breadth (Fig. S9-12; Table S4) both recovered the same relationships with caterpillar abundance that were found for taxonomic diet breadth. Importantly, more specialized species have greater local abundance, and that relationship is not sensitive to the measure of diet breadth used.

## Discussion

As a theoretical expectation, the “jack of all trades is a master of none” paradigm has persisted despite decades of examination with limited consensus, depending on the organisms and the traits being examined (Futuyma & Moreno 1988; Forister *et al.* 2012). For many morphological traits, biophysical constraints in vertebrates have produced clear examples where an ecological generalist underperforms specialists evaluated in a common context or on a shared resource (Oufiero *et al.* 2012). The story has been more complicated for herbivorous insects, where the focal trait is typically developmental performance in juvenile life history stages, and specialists have often not outperformed generalists when experimentally evaluated on a common resource (García-Robledo & Horvitz 2012; Agrawal 2020; Hardy *et al.* 2020). In the absence of a single unifying mechanism explaining the causes of ecological specialization, our results confirm that species abundance patterns associated with diet breadth reveal emergent patterns at the community level. Instead of an experimental framework, we have taken a different approach by examining the relationship between abundance and diet breadth measured across large spatial and temporal scales in the field, and we find that specialists tend to be locally more abundant while generalists occupy more places on the landscape.

Despite variation in the strength and direction of the relationship between diet breadth and abundance across different levels of observation and measures of abundance we reported here, the raw effect sizes were substantial. At the smallest spatial scale (plot-level), adding an additional host plant species to taxonomic diet breadth results in a 4% decrease in caterpillar density per available host plant (Fig. 2A, Table 1, Fig. S1). This result raises the possibility that specialists are indeed better adapted to their host plants, while generalists benefit from being able to colonize more hosts. That general pattern is largely robust to observations taken at different elevational and spatial scales (Fig. 2), although the signal was quite weak at the very largest scale that included general collecting (Fig. 3), possibly because our measure of “abundance” at that level is not as sensitive as the plot-level counts of individuals used at the smaller spatial scales. The positive relationship between diet breadth and occupancy is an interesting contrast to the smaller scale negative relationship, but the patterns are not inconsistent and reflects high turnover of specialist caterpillars across the landscape and low turnover of generalists. In fact, a null model predicts a strong, positive relationship between diet breadth and occupancy across the landscape (Forister & Jenkins 2017).

It is important to note that the results reported here do not address mechanisms generating the observed relationships between diet breadth and abundance or occupancy. As discussed above, the idea of genetic trade-offs as an expectation associated with the jack-of-all trades hypothesis (Levins 1968; Agosta & Klemens 2009) has a long history with only minimal support for herbivorous insects (Gompert & Messina 2016). Other equally-powerful ideas have received less attention, including the hypothesis that narrow niches are associated with more efficient selection for beneficial alleles, and generalists are less well adapted to any one resource because selection is effectively diluted across multiple environments (Whitlock 1996). Regardless of the mechanism or mechanisms explaining the tendency towards higher local densities of more specialized species, an important finding from our work is not only the sign and magnitude of that relationship, but also the shape of the variance. In particular, the range of densities observed for specialists is orders of magnitude greater than the range for densities of generalists (Fig. 2A). This broad distribution of densities for specialists counterintuitively raises the possibility that being a host specialist is a more multifaceted condition than host generalization. Specialists, for example, likely contend with a different or more variable suite of natural enemies that affect local abundance in ways that are heterogenous and warrant further study (Singer & Stireman 2005). It would be interesting to examine differences in the ratio of specialists to generalists among host genera (Fig. S13) with an eye towards measuring the effects of natural enemies and plant chemistry on those ratios. Understanding shifts in specialist to generalist ratios across different scales of observation for selected host taxa could provide considerable insight into the causes and consequences of the dominance of specialist herbivores (Novotny *et al.* 2002, 2012; Forister & Jenkins 2017).

The diet breadth of rare species, defined as those that were found as single individuals (“singletons”) in extensive sampling, is complicated and poses particular challenges. Rare species in tropical forests represent a high portion of insect herbivores, contributing to high species richness in tropical insect communities (Price *et al.* 1995), and singleton records are made of both specialist and generalist herbivores (Novotný & Basset 2000). Species are rare for various reasons, including sampling bias and host specificity (Novotný & Basset 2000). For instance, there may be insufficient spatial and seasonal replication of sampling, as certain insect species may occupy the same host plants in different seasons. Insect herbivores depend on the quality of their host plants, and their preferred host plants may not be sampled, limiting the possibility of knowing their diet breadth. We removed over 1000 singletons from the analyses in the present study because they were found once in the 17 years of data collection. The data was collected all year-round; therefore, it is unlikely that the collection does not cover seasonal and spatial variation.

In conclusion, the results reported here confirm the long-standing, but rarely observed assumption of a negative correlation between abundance and diet breadth and are a reminder that many decades are often needed for the evaluation of basic assumptions that underlie ecological theory, especially when that theory is applied to complex traits such as digestion and juvenile development. It is also the case that such relationships are likely dependent on the taxa and communities being studied. A similar investigation was recently conducted with field data for armored scale insects, with the conclusion that diet breadth and abundance are not linked (Peterson *et al.* 2020). Although Peterson *et al.* (2020) did not detect an overall abundance-diet breadth relationship, it is the case that the most abundant species tended to be extreme specialists (consistent with the high variance we observed in our specialized species). There is clearly much yet to be learned about ecological and dietary specialization, and the observations reported here are made more relevant in the midst of ongoing reports of declining insect abundances from around the world (Salcido *et al.* 2020; Wagner 2020; Forister *et al.* 2021; Halsch *et al.* 2021). We hope that coming decades see long term datasets, such as the one used here, being used simultaneously to motivate conservation and to test long-standing and unresolved ideas about how the natural world works.

## Acknowledgements

We would like to thank the Ministry of the Environment of Ecuador for providing permits under the genetic access contract MAE-DNB-CM-2016-0045 and the project “Interacciones entre plantas, orugas, y parasitoides de los Andes del Ecuador.” from the Instituto Nacional de Biodiversidad - Ecuador. Funding was provided from Earthwatch Institute and the National Science Foundation (DEB 1442103). For substantial contributions to the field work and development of ideas discussed here, we thank countless community science volunteers, the UNR Plant-Insect Group, G.L. Gentry, J.O. Stireman, J. Miller, L. Richards, A. Smilanich, A. Glassmire, H.F. Greeney, L. Salagaje, and W. Simbaña.

