## Supplemental Figures and Tables for "Jack-of-all-trades paradigm meets long-term data: generalist herbivores are more widespread and locally less abundant"

!                                                                                                                                  "

"!

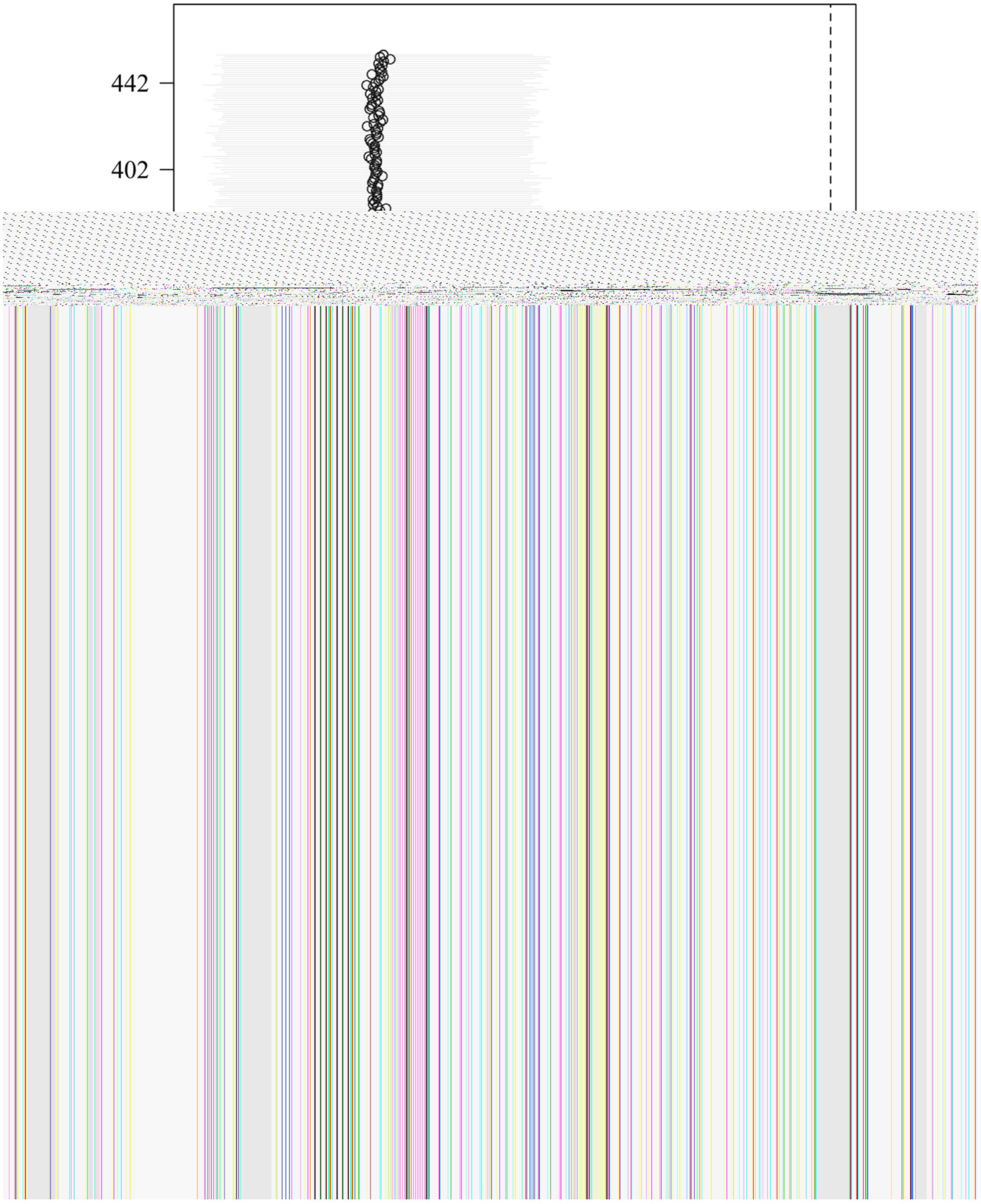


#!                                                                                                                                  !

8! !$ %&() *%+,) ,%-&*+!. %/,&0!1%2!3%+!' 2%!1%2!. ,%&42%. &5!' &6' 2,*7/ !13*&

A! ) 37/ &2!/ ,9%!:4*&5!,- !3*; 10!/ ) ' 3%!' - . !<=> !) 2%,43%- &26' 3!,/ !,- ) 37. % !,- !; 2%0!4' 2/?@

=! !

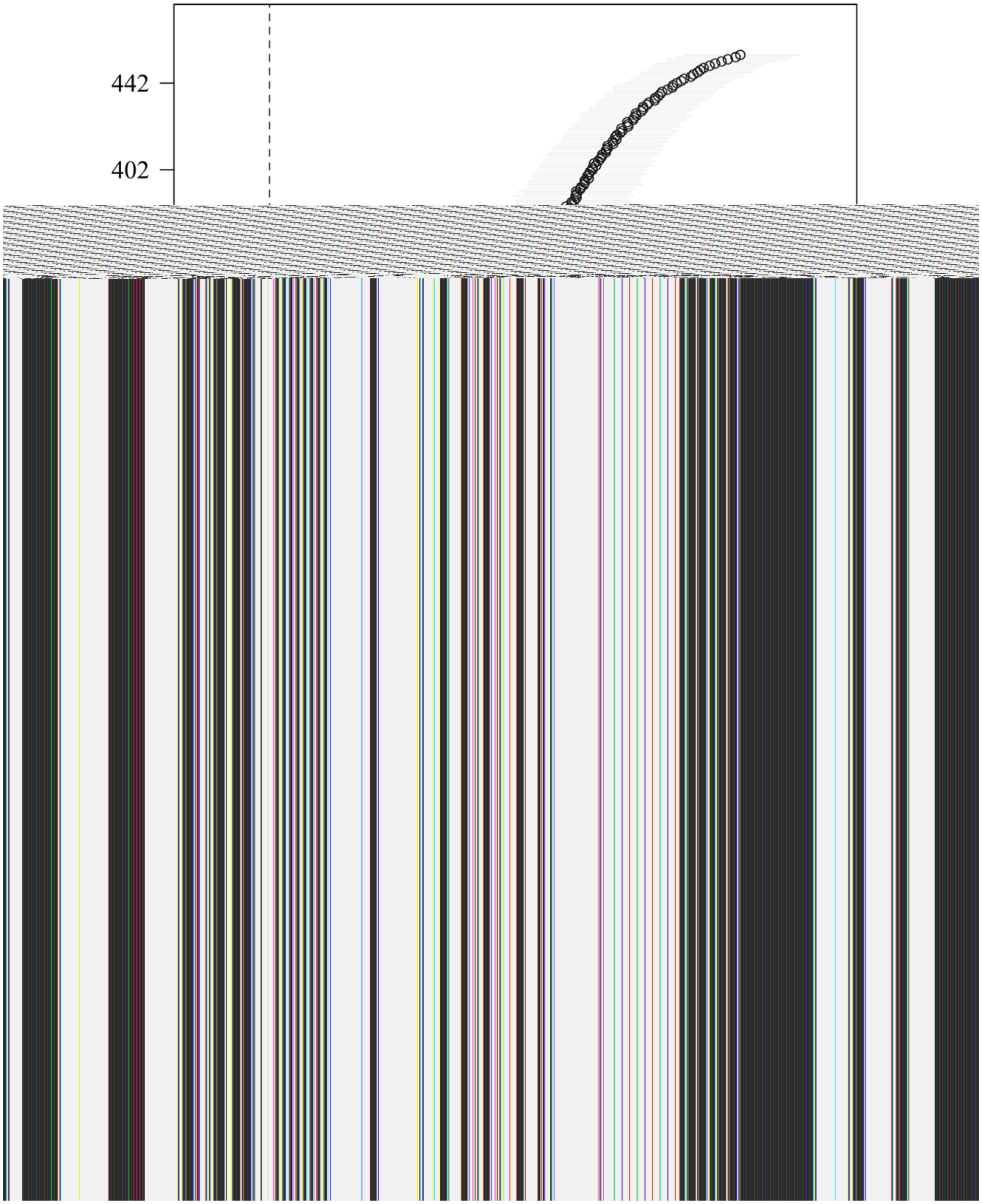


B! !

C! !$%&()*%+,),%-&*+!*))71'-)0!1%2!..,%&42%..&5!'&6'2,*7/!13*&)37/&2!

D! /,9%!:4*&5!,-!3*;10!/)'3%!'-.!<=>!)2%,43%,-&26'3!,/!,-)37.%!,-!;2%0!4'2?@!

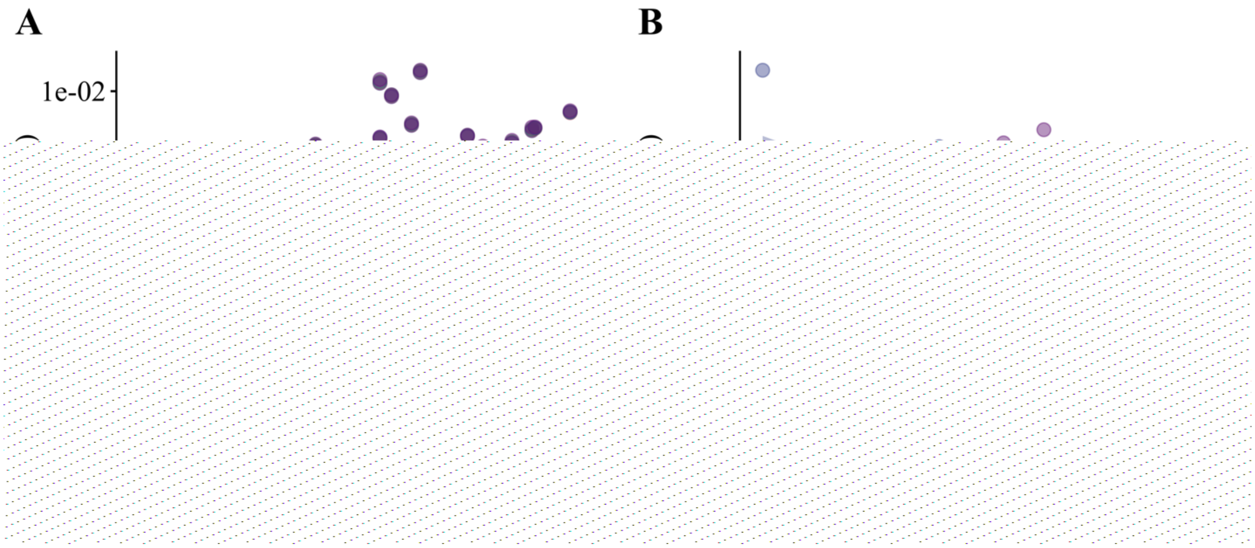


!F%3 &*-/5,1!" G*-; !. ,%&42%. &5!" -. !)' &21,33 2!. %/,&0!1%2!13*& 2%!" &

. ,+%2% &/)' 3%!*+!*4/%26' &*- H! ?!13*&/)' 3%1!" -. !4?!%3%6' &*- !/)' 3%@0 3*&3%6%3!. ' &!K%2%

)*G4,- % !+*2!%)5!)' &21,33 2!/1%),%!:1*,- &?!+*2!6' 2*7/!13*& ; ; 2%' &%!:437%H=!13*&!2% H#=I!

; 2%H=E!" -. !17213%#=E?!&!%6' 37' &%5%%+%)&*+!/)' 3%@ *2!5%%3%6' &*- !" -' 30/,/I!. ,/)2%&%

%3%6' &*- !4' -. /!,-)37. % H!E!(!" EEE!G!:2% ?I!" EE"(#EEE!G!:; 2% ?I!#EE"(8EEE!G!:17213%I!" -. !

' 4*6%8EEE!G!:437%@ *,- &!K%2%/%&!" !& -/1' 2%)0!6' 37%; ,6%!5%M' -&0!*+!1*,- &I!/7)5!

&' & ' 2N%2!57%!,-. ,)' &%!5,; 5!. %/,&0!*+!1*,- &@ !

!

!

!

!

!

!

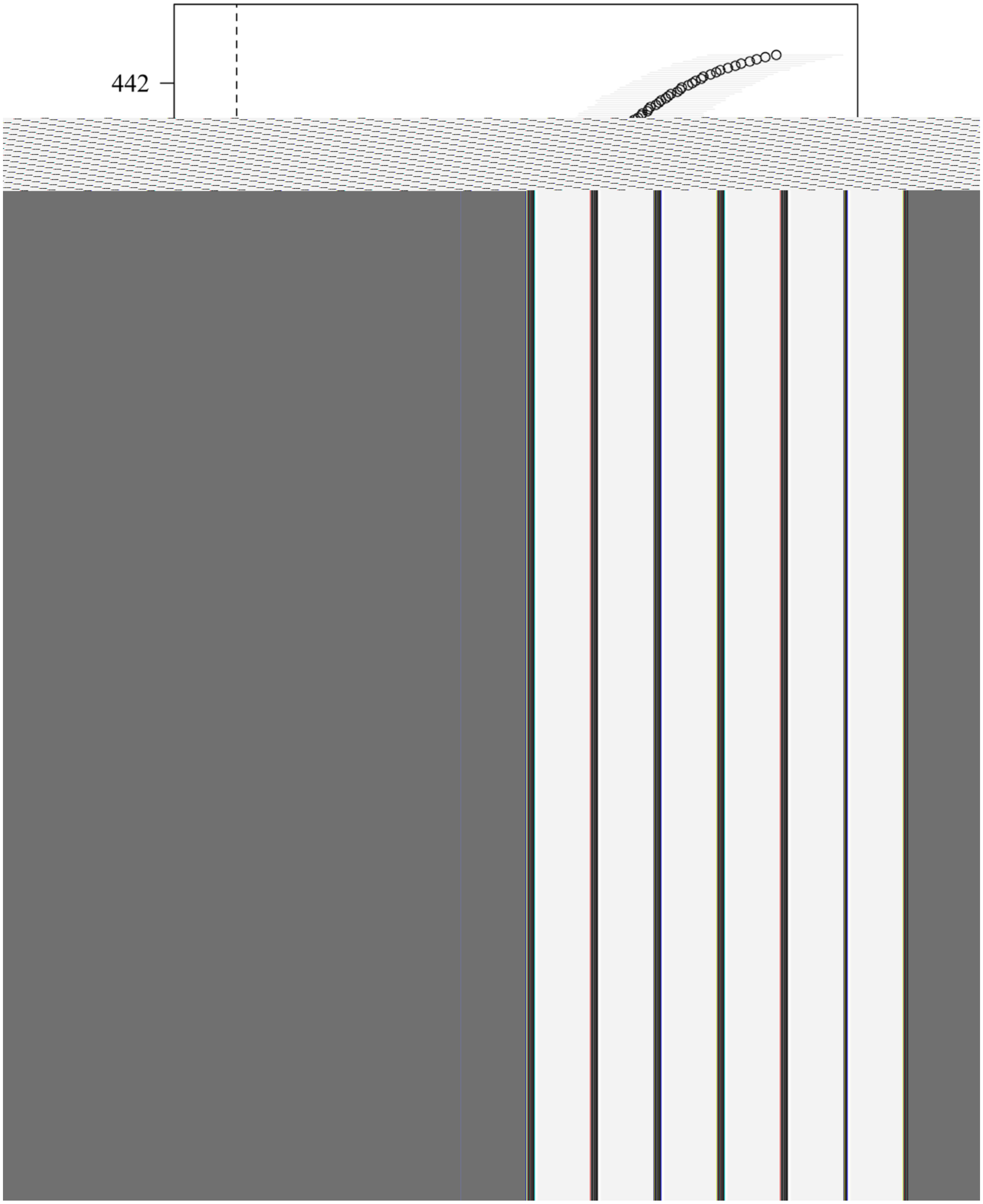


8#! !

88! !$ %&() *%+,) ,% &*+!*)) 71' - ) 0!1%2!*2 ,-' &% !. ,%&42%. &5!' &6' 2,*7/ !13*&

8A! )37/ &22!/ ,9%!:. %/ ,&0!1%2!3%+!' 2%!,/ !,- !3*; 10!/ )' 3%!47&- *&&5%1*2 ,-' &% !. ,%&42%. &5@!O5%<=> !

8=! )2%,43%,- &26' 3!,/ !,- )37. %!,- !; 2%0!4' 2/ ?@

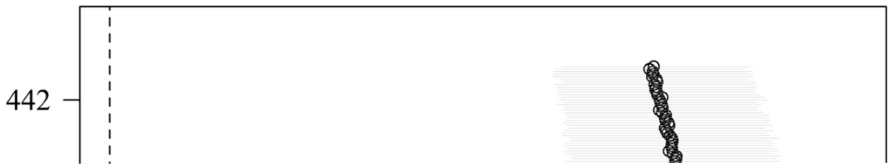


8B! !

8C! [illegible]

8D! [illegible]

8<! [illegible]

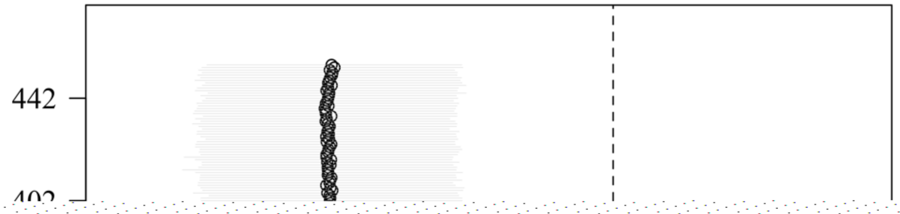


A=! !

AB! !$ %&() *%+,) ,% &*+!. %/ ,&0!1%2!3%+!' 2% !6/ @9(/) *2%*+!1503*; % %&) !. ,%&

AC! 42%. &5!' &6' 2,*7/ !13*&) 37/ &22!/ ,9%!:4*&5!,- !3*; 10!/) ' 3%!' - . !<=> !) 2% ,43%,- &26' 3!,/ !,- ) 37. % !,- !

AD! ; 2%0!4' 2/ ?@

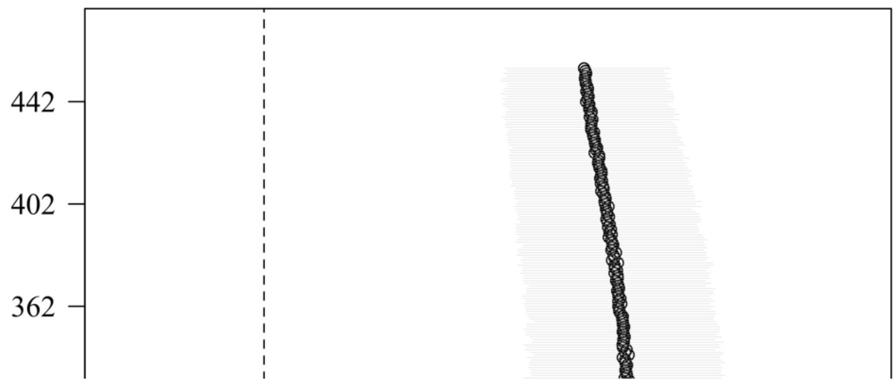


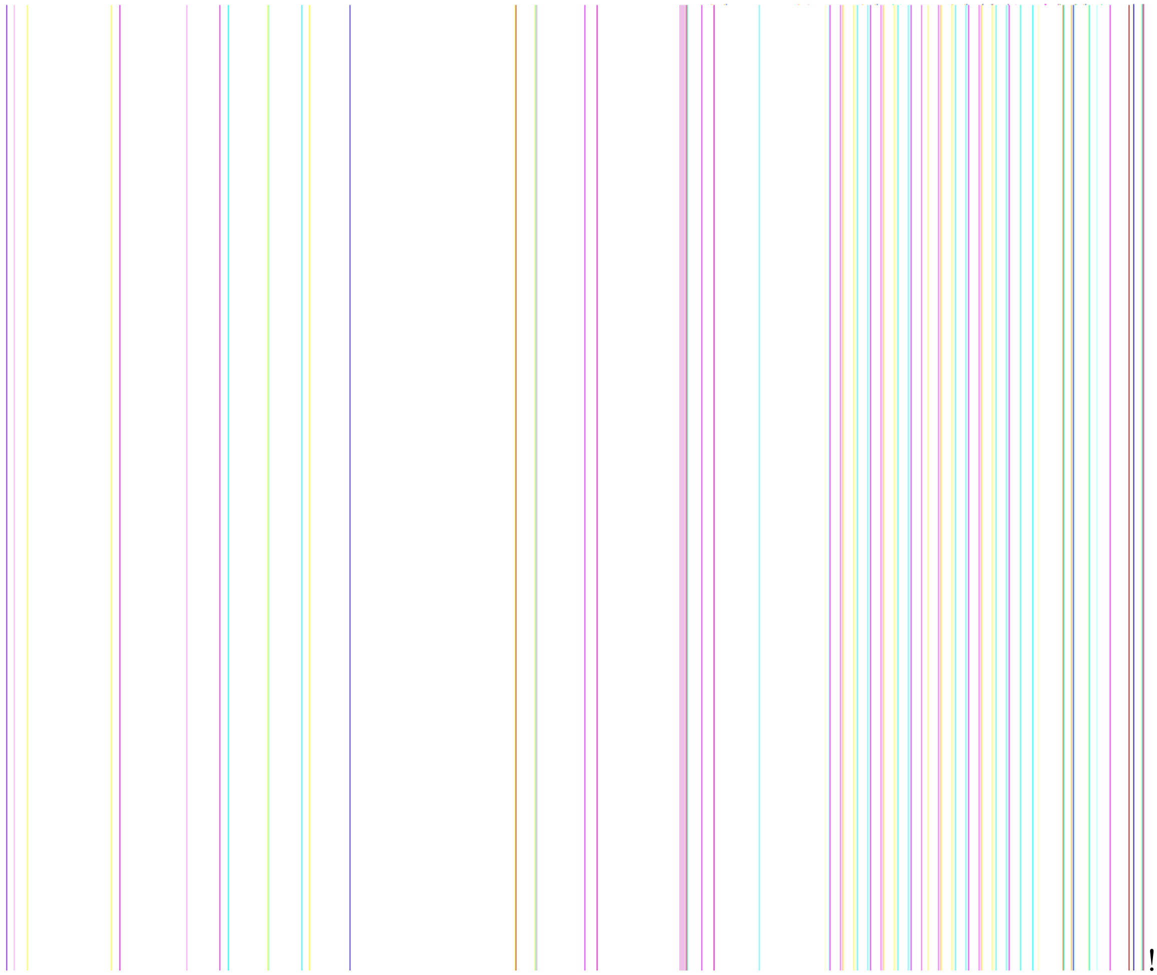

=8! !

=A! !$ %&() *%+,) ,% &*+!. % / ,&0!1%2!13*& 2% !6/ @9(/) *2%*+!1503*; % %&) !. ,%&

==! 42%. &5!' &6' 2,*7/ !13*&) 37/ &22!/ ,9%!:4*&5!,- !3*; 10!/) ' 3%!' - . !<=> !) 2% ,43%,- &26' 3!,/ !,- ) 37. % !,- !

=B! ; 2%0!4' 2/ ?@

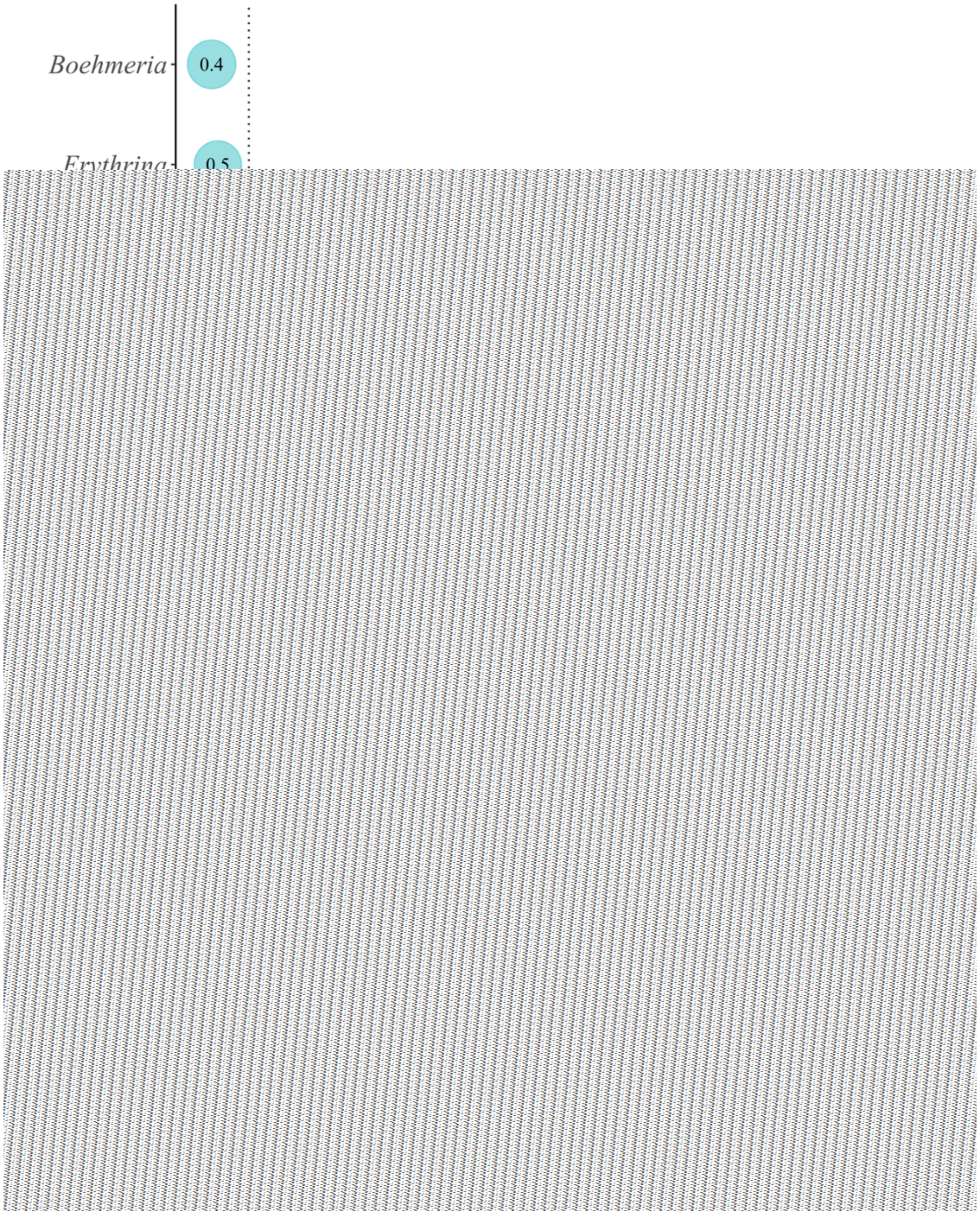


O5%2' &*!*+!/1%),' 3/&&*!; %%2' 3/&)' &&21,33' 2/!*-!/5' 2%!5*/&13 - &

; %%2' !6' 2,%@U' 37%!K,&,-!1*,-&!,-. ,)' &1&%/1%),' 3/&&*!; %%2' 3/&2' &*!+*2!&%1G*/&

)*GG*-30!/' G13%!13 - &; %%2' !K,&,-!13*&@V,9%!*+!&%1*,-&' 2%12*1*2&*-' &1&*!&%

' 6' ,3 43%5*/&13 - &' 2%!%&G' &%!' )2*//!' 33/' G13%!13*&!+*2!K5,)5!)' &21,33' 2/!K%2%+*7-. @

Q*3*2/!. ,+%2%&' &1; 2%&22!. %/,&0!*+!; %%2' 3/&!:437%3!*2!/1%),' 3/&!:2%?@

**Supplemental Table 1**. Summary of different measures of ecological specialization.

| ! | ! | "#$%&()! | *(+&$, (-)! | . &0$&)&! | *&1&(&2!3 &4&+&%&! |
|---|---|---|---|---|---|
| **5 &)6+&!#4!** | | | | | |
| **&%1#, 7%1!** | | | | | *et al.* |
| **)' &71 18/ (7#$** | | | | | *et al.* |
| | | | | | *et al.* |
| | | | | | *et al.* |
| | | | | | *et al.* |

*et al.*

*et al.*

**5 &)6+&)!#4!** *et al.*

**9 7&(!: +&2(- !**

*et al.*

*et al.*

*et al.*

*et al.*

*et al.*

73

**Supplemental Table 2.** Point estimates for beta coefficients (bold) and associated 95% credible intervals for relationship between taxonomic diet breadth and density per available plot area at plot and elevation levels. Plot levels include different cluster sizes which represent the number of aggregated 10-m plots.

| **"#$#%&!( ) *#+$, -.&!** | **0 #/ *.-1!2#+!, $, .%) %!2 %&!, +#, !** |
|---|---|
| **3 %&!%$#%5%*-#+!*.7#8!** | |
| 5 | **9:; <!**[0.52, 0.77]**!** |
| 25 | **9:; ; !**[0.54, 0.78]**!** |
| 50 | **9:; =!**[0.55, 0.83]**!** |
| 250 | **9:>?!**[0.59, 0.88]**!** |
| **@%$, -.&!%$#%** | |
| 0 m - 1000 m | **A9:9B<!**[-0.77, 0.73]**!** |
| 1001 m - 2000 m | **9:C>!**[0.24, 0.51]**!** |
| 2001 m - 3000 m | **9:; B!**[0.49, 0.75]**!** |
| > 3000 m | **AC:BB!**[-0.93, 0.27]**!** |

**Supplemental Table 3.** Point estimates for beta coefficients (bold) and associated 95% credible intervals for relationship between family-level ordinated diet breadth and different abundance indices at plot and elevation levels. Plot levels include different cluster sizes which represent the number of aggregated 10-m plots.

| [illegible] | [illegible] | [illegible] | [illegible] |
|---|---|---|---|
| **[illegible]** | | | |
| 5 | **[illegible]** [-3.30, -1.18] | **[illegible]** [3.94, 5.44] | **[illegible]** [3.43, 5.64] |
| 25 | **[illegible]** [-3.42, -1.21] | **[illegible]** [3.06, 4.44] | **[illegible]** [3.57, 5.84] |
| 50 | **[illegible]** [-3.35, -1.20] | **[illegible]** [2.50, 3.86] | **[illegible]** [3.71, 6.03] |
| 250 | **[illegible]** [-3.26, -1.12] | **[illegible]** [0.68, 2.17] | **[illegible]** [3.91, 6.62] |
| **[illegible]** | | | |
| 0 m - 1000 m | 38.66 [9.03, 60.12] | **[illegible]** [-14.90, 11.75] | **[illegible]** [-24.44, 15.08] |
| 1001 m - 2000 m | **[illegible]** [-2.69, -0.10] | **[illegible]** [1.89, 3.39] | **[illegible]** [1.09, 3.18] |
| 2001 m - 3000 m | **[illegible]** [-3.67, -1.42] | **[illegible]** [3.86, 5.51] | **[illegible]** [3.19, 5.41] |
| > 3000 m | **[illegible]** [-11.37, 3.89] | **[illegible]** [-3.05, 1.19] | **[illegible]** [-8.60, 2.34] |

**Supplemental Table 4.** Point estimates for beta coefficients (bold) and associated 95% credible intervals for relationship between family-level phylogenetic diet breadth and different abundance indices at plot and elevation levels. Plot levels include different cluster sizes which represent the number of aggregated 10-m plots.

| **"#$#%&!( ) *#+$, -.& !** | **0 #/ *.-1!2#+!, $, .%) %! D&*-!2 %/ -!, +#, !** | **( 5562, / 51!** | **0 #/ *.-1!2#+!, $, .%) %! 2 %&!, +#, !** |
|---|---|---|---|
| **3 %&!%$#%5%*-#+!*.7#8!** | ! | ! | ! |
| 5 | **A9:BB** [-0.16, -0.061] | **9:??** [0.19, 0.26] | **9:?B!** [0.16, 0.27]! |
| 25 | **A9:BB** [-0.17, -0.063] | **9:B=** [0.15, 0.22] | **9:??!** [0.16, 0.27]! |
| 50 | **A9:BB** [-0.17, -0.062] | **9:B<** [0.12, 0.19] | **9:?C!** [0.17, 0.28]! |
| 250 | **A9:BB** [-0.16, 0.059] | **9:9>** [0.037, 0.11] | **9:?<!** [0.18, 0.32]! |
| **@%$, -.& !%$#%** | | | |
| 0 m - 1000 m | **?:<E** [0.83, 3.95] | **A9:9; E** [-0.79, 0.86] | **A9:CF!** [-1.73, 0.75]! |
| 1001 m - 2000 m | **A9:9=E** [-0.16, -0.0055] | **9:BE** [0.094, -0.18] | **9:B9!** [0.042, 0.17]! |
| 2001 m - 3000 m | **A9:BE** [-0.20, -0.081] | **9:?<** [0.20, 0.29] | **9:?C!** [0.16, 0.29]! |
| > 3000 m | **A9:?9** [-0.62, 0.23] | **9:9<9** [-0.16, 0.068] | **A9:B=!** [-0.45, 0.11]! |
